# Optimized semi-specific PCR amplification using arbitrarily degenerate primer for genome-wide genotyping and its application in peanut genetic diversity study

**DOI:** 10.1101/2024.08.04.606550

**Authors:** Sheng Zhao, Yue Wang, Xuejiao Zhang, Shuxian Xie, Haotian Chen, Yiming Yan, Jiaqi Gu, Peng Chen, Yuxiao Chang, Zhijun Xu

**Affiliations:** Guangdong Laboratory of Lingnan Modern Agriculture, Genome Analysis Laboratory of the Ministry of Agriculture and Rural Affairs, Agricultural Genomics Institute at Shenzhen, Chinese Academy of Agricultural Sciences, Shenzhen 518120, China; Guangdong Modern Agriculture (Cultivated Land Conservation and Water-Saving Agriculture) Industrial Technology Research and Development Center, Zhanjiang Experiment Station, Chinese Academy of Tropical Agricultural Sciences, Zhanjiang 524031, China

**Keywords:** whole genome genotyping, semi-specific amplification, peanut, germplasm resource, population structure

## Abstract

Cultivated peanut (*Arachis hypogaea* L.) is an important legume crop with a large genome size but a limited genetic diversity. Evaluating the genetic variation among diverse peanut germplasms using genome-wide molecular markers is an effective strategy to explore its genetic diversity and thereby facilitate peanut improvement. In this study, we introduced a novel whole-genome genotyping (WGG) technique named dRAPD-seq (degenerate Random Amplification Polymorphic DNA and sequencing), which relies on semi-specific PCR amplification by arbitrarily degenerate (AD) primer and next-generation sequencing (NGS), and demonstrated its robust reproducibility and high accuracy. Subsequently, we applied dRAPD-seq to investigate the genetic relatedness within a population of 101 diverse peanut accessions and identified a low genetic diversity among these accessions. Our phylogenetic tree, population structure analyses, and principal component analysis (PCA) indicated that this population could be clustered into three subpopulations, largely corresponding to three botanical types. In conclusion, this research not only introduced a cost-effective and easy-to-conduct WGG method but also provided valuable insights for utilizing these peanut accessions in future breeding programs.

## Introduction

The cultivated peanut (*Arachis hypogaea* L.) is a significant allotetraploid (AABB, 2n = 4×= 40; ~2.6 Gb) legume crop, providing essential edible oil and protein for human consumption. Since its origination from South America approximately 9400 years ago, the peanut has been extensively cultivated and distributed in over 100 countries across tropical and temperate regions worldwide [1]. Typically, cultivated peanuts are categorized into two subspecies, subsp. *hypogaea* and subsp. *fastigiata*, distinguished by the arrangement of floral axes on the main stem. These subspecies further divide into six botanical types: Viriginia, Hirsute and Peruviana in subsp. *hypogaea*, and Valancia, Spanish and Aequatoriana in subsp. *fastigiata* [2,3]. As with many crops, the effective characterization and evaluation of peanut germplasm are essential prerequisites for its efficient utilization in breeding and research endeavors [4]. China, being one of the world’s largest peanut producers and consumers, has greatly benefited from the utilization of diverse peanut germplasms collected domestically and internationally [5]. Through phenotypic selection, significant improvements have been achieved in yield, disease resistance, oleic acid composition, and other agronomic traits in peanut breeding [6]. However, Chinese peanut cultivars exhibit a limited genetic diversity due to the frequent use of closely related genetic resources, with pedigree analysis confirming that over 70% of cultivars have common ancestors ‘Fuhuasheng’ and ‘Shitouqi’ [7]. Despite the availability of numerous molecular markers, less than 15% of the entire peanut collection has been genotyped at the molecular level so far, mainly because of the laborious and expensive genotyping process [8–10]. To broaden the genetic variation of cultivated peanut in future breeding program, it’s critical to develop a cost-effective and high-throughput genotyping technology to assess the genetic diversity and population structure among the diverse peanut germplasm resources that have been widely released.

DNA-based molecular markers are widely acknowledged as the most reliable tools for assessing genetic variability and diversity in plants. Despite the successful use of various molecular markers such as restriction fragment length polymorphism (RFLP), random amplification of polymorphic DNA (RAPD), and amplified fragment length polymorphism (AFLP) in exploring genetic diversity among different peanut germplasms [11–13], simple sequence repeat (SSR) and single nucleotide polymorphism (SNP) are generally preferred due to their co-dominant nature, high-throughput capability, and ease of use [2,14,15]. In recent years, SNP-based genotyping technology has gradually supplanted SSR in peanut genetic diversity researches [5,9,16,17]. This shift is primarily attributed to significant advancements in NGS technologies and the improved availability of high-quality peanut genome assemblies [1,18,19].

Library preparation is an indispensable part of the NGS. Regardless of the sequencing platform or technology used, a typical library preparation procedure consists of two main steps. Initially, complete genomic DNA (gDNA) is fragmented into short fragments (typically 250–600 bp) through various methods, such as sonication or transposase-mediated gDNA breakage for whole genome sequencing (WGS) [20], or restriction endonuclease-mediated DNA digestion for Genotyping-by-Sequencing (GBS) [21–23]. Subsequently, adapters specific to the sample are ligated to both ends of the short fragments using techniques like sticky-end ligation, complementary A-T base pairing, or PCR amplification. To address the challenges associated with specialized instruments, costly reagents, or laborious processes required for gDNA fragmentation, several specific PCR-based genotyping technologies have been developed recently, such as target SNP-seq [24], multiplex restriction amplicon sequencing (MRASeq) [25], and Hyper-seq [26]. However, these new genotyping methods necessitate relatively complex primer design steps, potentially limiting their broad application across diverse crop species.

In our previous study, we accidentally discovered that any sequence specific primer (SP) possesses the intrinsic capability to anneal and prime a semi-specific amplification through primer-template mismatched annealing (PTMA) along the genome, resulting in the generation of tens of thousands of stable PCR products. Building upon this finding, we have developed two methods: foreground and background integrated genotyping by sequencing (FBI-seq) [27] and NGS-enhanced thermal asymmetric interlaced PCR sequencing (TAIL-peq) [28]. In addition, the arbitrary selection of a SP primer for RAPD has proven successful as a DNA molecular marker assay method [29], and further investigation into the relationships between primer sequences and amplification patterns revealed that the rare and fortuitous PTMA at both DNA ends eventually lead to the amplification of encompassing DNA segments [30,31]. However, it is worth noting that DNA smearing often occurs alongside clear bands in RAPD gel electrophoresis assays. Considering the abundance of PTMA events with SP primers when using NGS for PCR product analysis [27], we speculate that the smearing observed in RAPD gel electrophoresis assays may result from PTMA at multiple other loci. Additionally, an AD primer is likely to amplify more loci distributed throughout the genome compared to a SP primer used in the RAPD method. In this study, we presented a novel WGG method based on the semi-specific amplification by AD primer and NGS sequencing, and then utilized it to characterize the genetic diversity and population structure of a peanut germplasm resource population. Our results provide not only a novel cost-effective and easy-to-conduct genotyping method, but also valuable insights into the genetic relationships among peanut accessions, which could enhance their effective utilization for future peanut breeding efforts.

## Results

### The workflow of dRAPD-seq

The dRAPD-seq method comprised several key steps. Initially, in the preliminary semi-specific PCR reaction, AD primers ligated with unique barcodes were randomly bound to the gDNA through PTMA (Fig. 1A). Subsequently, PCR products from many samples were pooled into one tube by AIO-seq strategy [32], since that amplicons corresponding to each sample could be distinguishable by distinct barcodes (Fig. 1B). The pooled PCR products then underwent an A-tailing reaction (Fig. 1C), followed by the ligation of Y-shaped Illumina adapters containing a T-base overhang and integrated within the index sequence, with DNA fragments via A-T base pairing (Fig. 1D). After that, the resulting TruSeq library was prepared for subsequent NGS sequencing (Fig. 1E). During the sequencing process, Read1 and Read2 sequences were generated, commencing with distinct barcodes (Fig. 1F). Throughout the whole workflow of dRAPD-seq, only one AD primer and a PCR reaction were involved, thus could be regard as one of the simplest high-throughput genotyping methods.

**Figure 1.**
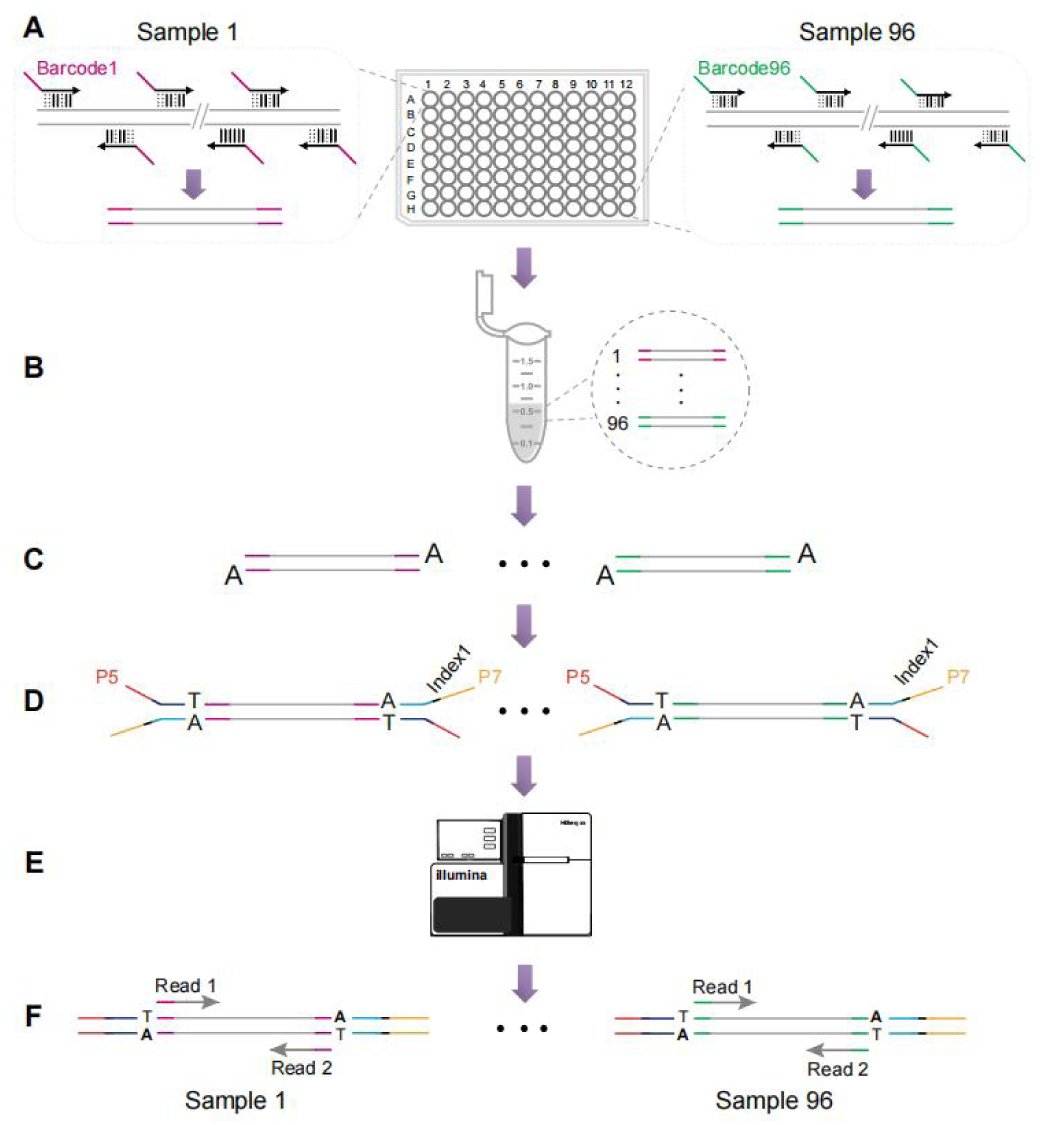
Graphical representation of dRAPD-seq (degenerate Random Amplification Polymorphism DNA and sequencing) workflow. **A**, same arbitrary degenerate (AD) primer but ligated with distinct barcodes were binding to the genomic DNA via primer-template mismatched annealing (PTMA). **B**, barcoded amplicons of all samples were pooled into one tube. **C**, an A-base is added to the blunt ends of pooled PCR products. **D**, Y-shaped Illumina adapter sequences (embedded within the index) were ligated to the fragments using A-T base pairing. **E**, the resulting TruSeq library was subjected for high-throughput sequencing. **F**, Read1 and Read2 sequences were produced from DNA library.

### Feasibility and genome characterization of dRAPD-seq

To investigate the feasibility and characteristics of dRAPD-seq data, we randomly designed an AD primer (AD1_Bc1; Supplementary Data Table S1) to amplify the rice Nipponbare genome and performed three technical replicates. Except for some specific peaks, the whole products were appeared as a long smear after initial semi-specific PCR amplification (Supplementary Data Figure S1). Through NGS sequencing, we obtained ~5.4–6.8 Gb data for each of three replicates and then randomly subsampled 1.0 Gb reads from each replicate for subsequent comparative analysis. When aligning these data to the Nipponbare_v7.0 reference genome [33], we observed a consistent distribution of reads across the entire genome in all three replicates (Fig. 2 A).

**Figure 2.**
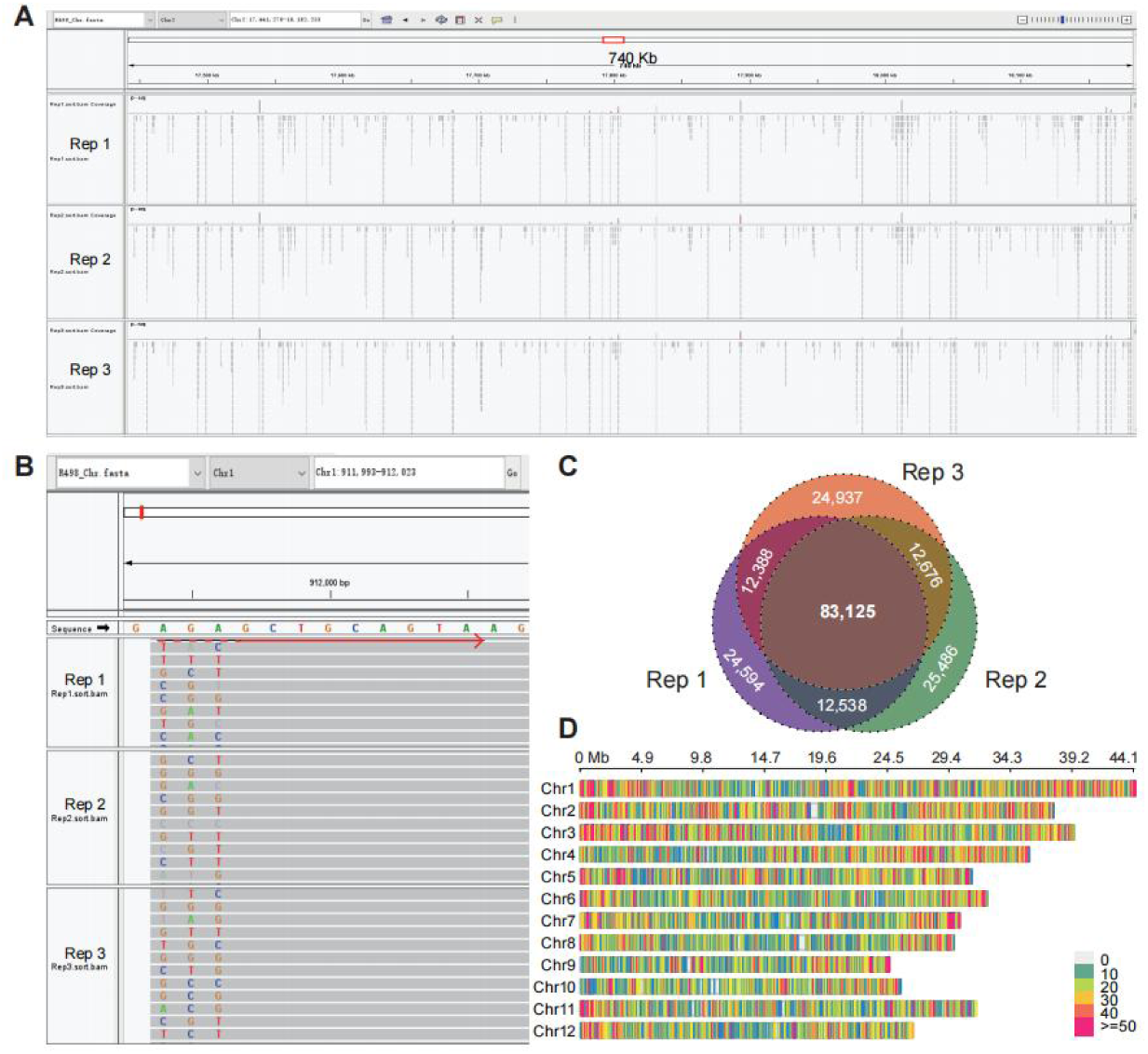
Genome characterization of the dRAPD-seq data. Three technical replicates were performed when using arbitrary degenerate (AD) primer (AD1_Bc1) for PCR amplification and DNA library preparation. **A**, a snapshot of the Integrative Genomics Viewer (IGV) browser displaying the genome distribution of dRAPD-seq reads from three replicates. **B**, a IGV snapshot showing that AD primer was bound with DNA template by primer-template mismatched annealing (PTMA) and dRAPD-seq reads were accumulated to form a tag. Red right arrow represented the binding site of AD primer. **C**, number of shared tags among three technical replicates. **D**, distribution of 83,125 shared tags across the rice genome. Distributions were calculated within a 0.1-Mb bin size.

Although the overall genome coverage was modest as 8.10–8.22% across the three replicates, we were able to identify a total of 132,645–133,825 genome-wide sequence tags (Supplementary Data Table S2), which were formed by PTMA (Fig. 2 B; Supplementary Data Figure S2). The mean coverage depth of these tags in the three replicates reached up to 29.73×, 29.90×, and 29.06×, respectively. Among these high-depth tags, nearly 83,125 tags, representing 62.11–62.67% of the all, were common to all three replicates (Fig. 2 C), and they were well-distributed across all the chromosomes of rice genome (Fig. 2 D). These results indicated that semi-specific amplification initialed by AD primer could consistently yield high-depth reads that were enriched at partial genomic regions in the dRAPD-seq method.

### Reproducibility and accuracy of dRAPD-seq method

To examine the reproducibility of dRAPD-seq reads, we first calculated the genome coverage at depths ranging from 1× to 50×, and found that corresponding values at the same depth were consistently comparable among the three replicates (Fig. 3 A). Subsequently, we measured the mean read depth and genome coverage within each 0.1 Mb sliding-window and both found a high level of consistency across the three replicates (Supplementary Data Figure S3), with Pearson correlation coefficients of 99.83 ± 0.05% (mean ± S.D.) and 95.18 ± 0.04%, respectively (Fig. 3 B and C). Finally, we drew Lorentz Curves to evaluate the read coverage uniformity throughout the genome, and discovered a high level of reproducibility among the three replicates, albeit with slight deviation from the diagonal line (Fig. 3 D). Taken together, these results underscored the robust replicability of the dRAPD-seq method.

**Figure 3.**
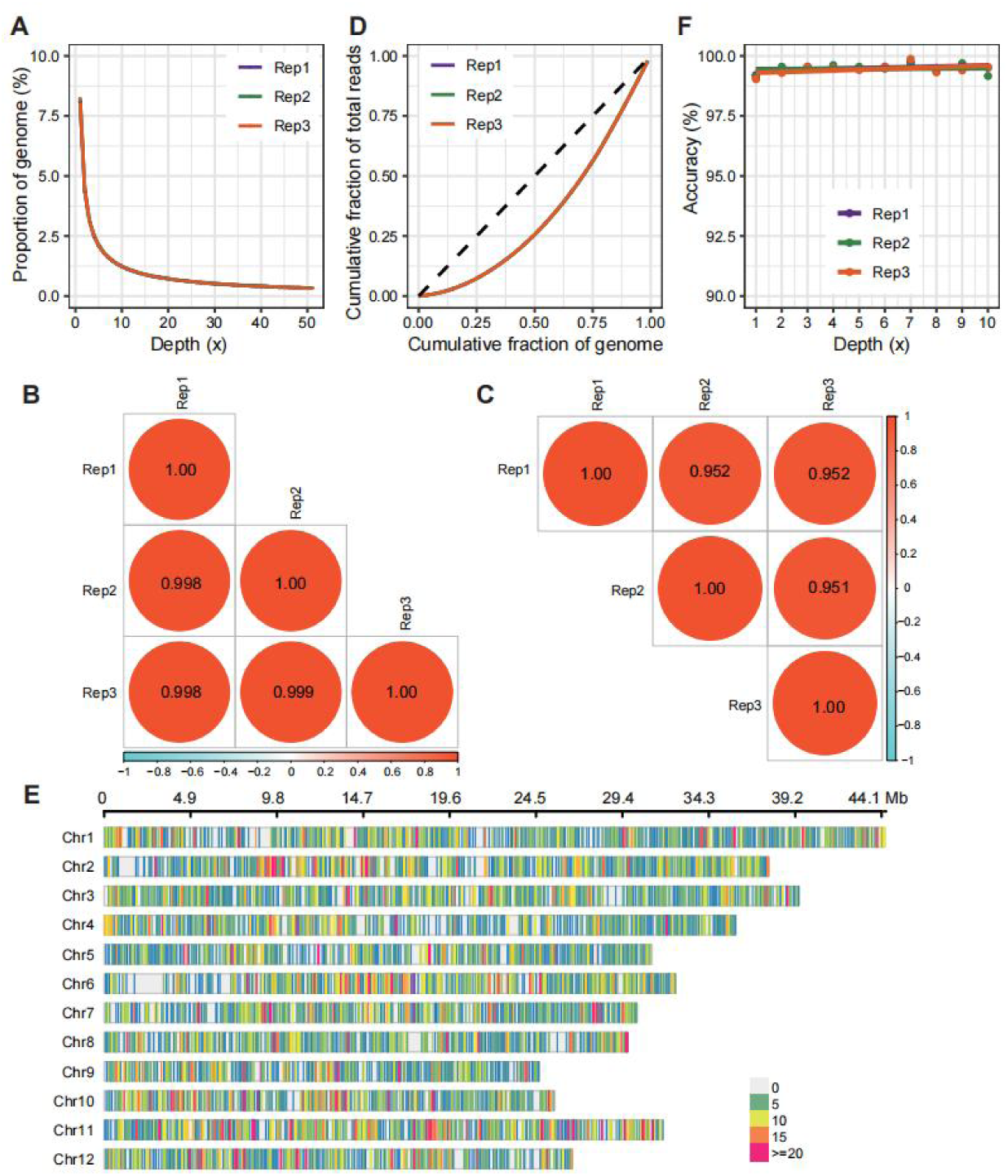
Reproducibility and accuracy of dRAPD-seq. **A**, evaluation of the breadth of coverage, defined as the percentage of base that has been sequenced for a given number of times (here 1× to 51×) in each of three technical replicates. **B**, Pearson correlation coefficient of mean read depth within a 0.1 Mb sliding-window calculated among three replicates. **C**, Pearson correlation coefficient of genome coverage within a 0.1 Mb sliding-window calculated among three replicates. **D**, lorenz curves of three technical replicates. A Lorenz curve illustrates the cumulated fraction of dRAPD-seq reads as a function of the cumulated fraction of the rice genome. The black dotted line represents perfectly uniform coverage. **E**, distribution of 23,759 shared SNPs across the rice genome. **F**, SNP accuracy calculated at varying coverage depth thresholds (1× to 10×) in each of three replicates.

To further assess the reliability and accuracy of dRAPD-seq genotyping method, we aligned the abovementioned 1.0 Gb of sequencing data from each replicate to the rice R498 reference genome [34], and detected a total of 53,345–54,415 SNPs across the three replicates (Supplementary Data Table S2). Notably, 23,759 SNPs (approximately 43.66–44.54% of the total) were identified as common among all replicates, showing genome-wide and uniform distribution throughout the rice genome (Fig. 3 E). In parallel, we performed the WGS on the Nipponbare gDNA and obtained 25.63 Gb of sequencing data (~ 65.71× depth), leading to the identification of 2,287,783 high-quality SNPs. We thus checked the SNP accuracy of dRAPD-seq method using WGS data served as gold standard, and found an increasing accuracy with higher SNP depth, reaching 99.21%, 99.14%, and 99.03% for the three replicates even at a depth as low as 1× (Fig. 3 F). Collectively, these results highlighted the robustness and high accuracy of the dRAPD-seq genotyping method.

### Genetic variations and fingerprint of peanut accessions

After the de-multiplexing of raw dRAPD-seq data, sequencing reads ranging from 0.53 to 21.61 Gb were obtained for the 101 peanut accessions, corresponding to genome coverages of 0.30–2.04% (Supplementary Data Table S3). On average, approximately 4.74 ± 3.45 Gb of sequencing reads and 0.98 ± 0.37% genome coverages were observed. Despite we found a positive correlation (Pearson *r* = 0.64, *p* < 0.001), the genome coverages remained low at 2.04% even at a sequencing depth of 10.12× (Fig. 4 A), indicating a notable enrichment of dRAPD-seq reads on a small subset of peanut genome. By applying stringent filtering criteria, we identified a total of 1,203 high-quality SNPs in this population, with SNP counts ranging from 27 on Chr7 to 106 on Chr20 (Fig. 4 B; Supplementary Data Table S4). Furthermore, these SNPs were evenly distributed across the peanut genome, with a mean coverage depth of 48.98 ± 30.93× (Fig. 4-C; Supplementary Data Table S5).

**Figure 4.**
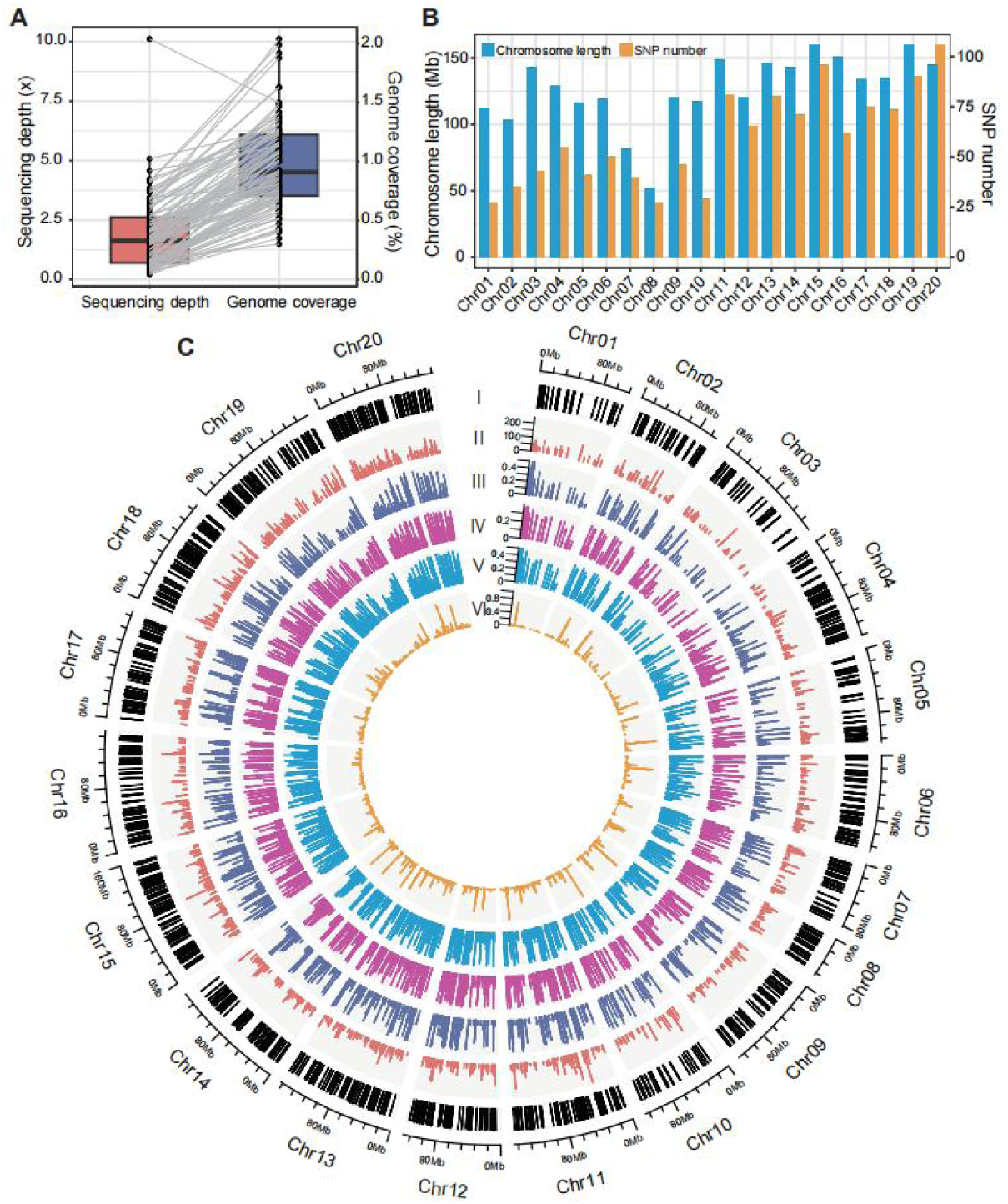
Sequencing and genetic characterization of 101 peanut accessions. **A**, relationship between the sequencing depth and genome coverage of each accession. Boxplots span from the first until the third quartile of the data distribution, and the horizontal line indicates the median value. The whiskers extend from the quartiles until the last data point within 1.5 times the interquartile range, with outliers beyond. Individual data points are also represented. Gray lines join samples pertaining to the same accession. **B**, SNP number on each of 20 chromosomes of peanut. **C**, circos plot displaying the genomic position (track I), mean coverage depth (track II), minor allele frequency (MAF, track III), polymorphic information content value (PIC, track IV), genetic diversity (Simpson’s Index, track V), and observed heterozygosity (*Ho*, track VI) of 1203 SNP markers.

The set of 1,203 high-quality SNPs identified in the 101 peanut accessions (Fig. 4 C; Supplementary Data Table S5) exhibited polymorphic information content (PIC) values ranging from 0.09 to 0.38, with an average of 0.23. Additionally, the mean minor allele frequency (MAF) was 0.20, ranging from 0.05 to 0.50, and the observed heterozygosity (*Ho*) averaged 0.08. Although *Ho* values varied widely from 0.0 to 1.0 across these SNPs, only 5.07% of the SNPs displayed *Ho* values exceeding 0.50. The genetic diversity coefficients (Simpson’s Index) ranged from 0.10 to 0.50, with an average of 0.28, and genetic distances spanned from 0.0218 to 0.464, averaging at 0.255 (Supplementary Data Table S6). Overall, these results suggested a low diversity and high genetic similarity within this peanut germplasm population. Furthermore, we generated DNA fingerprint profiles for the 101 peanut accessions using these refined SNPs (Supplementary Data Figure S4), offering potential molecular markers for peanut accession identification.

### Population structure analysis of peanut accessions

Using the high-quality SNPs, multiple analyses were conducted to explore the genetic relationships among 101 peanut accessions. Initially, we conducted phylogenetic and population structure analyses, revealing the classification of these accessions into three main subpopulations (K = 3) designated as G1, G2, and G3, comprising 55, 10, and 36 accessions, respectively (Fig. 5 A and B; Supplementary Data Table S3). Interestingly, these subpopulations corresponded primarily to three distinct botanical types. In the G1 subpopulation, the majority of accessions (38 out of 55) belonged to the Spanish type (subsp. *fastigiata*), originating mainly from the Pearl River basin encompassing Guizhou, Guangdong, Guangxi, and Fujian provinces. The G2 subpopulation consisted predominantly of Valencia type accessions (subsp. *fastigiata*), with nearly all accessions (9 out of 10) originating from the northeast and north of China, particularly from Heilongjiang, Jilin, Hebei, and Shandong provinces. While the G3 subpopulation exhibited a variety of botanical types, primarily including 17 Virginia type (subsp. *hypogaea*) accessions, 7 Valencia type (subsp. *fastigiata*) accessions, and 4 intermediate type accessions. Furthermore, a broad geographic distribution of accessions was observed in this subpopulation, spanning the Yellow River basin (Shandong, Henan, and Hebei provinces) and the Pearl River basin (Guizhou, Guangdong, and Yunnan provinces). Additionally, there were 3, 3, and 5 foreign accessions classified in the G1, G2, and G3 subpopulations, respectively. Subsequently, we performed PCA analysis on this population, confirming the presence of the three distinct groupings aligning with the subpopulation lineages G1-G3 (Fig. 5 C), in accordance with the findings of the phylogenetic and admixture analyses. The first two principal components explained 37.25% of the total variance, with PC1 reflecting the variability of G1 and G2 groups and PC2 distinguishing G3 accessions from G1 and G2 samples.

**Figure 5.**
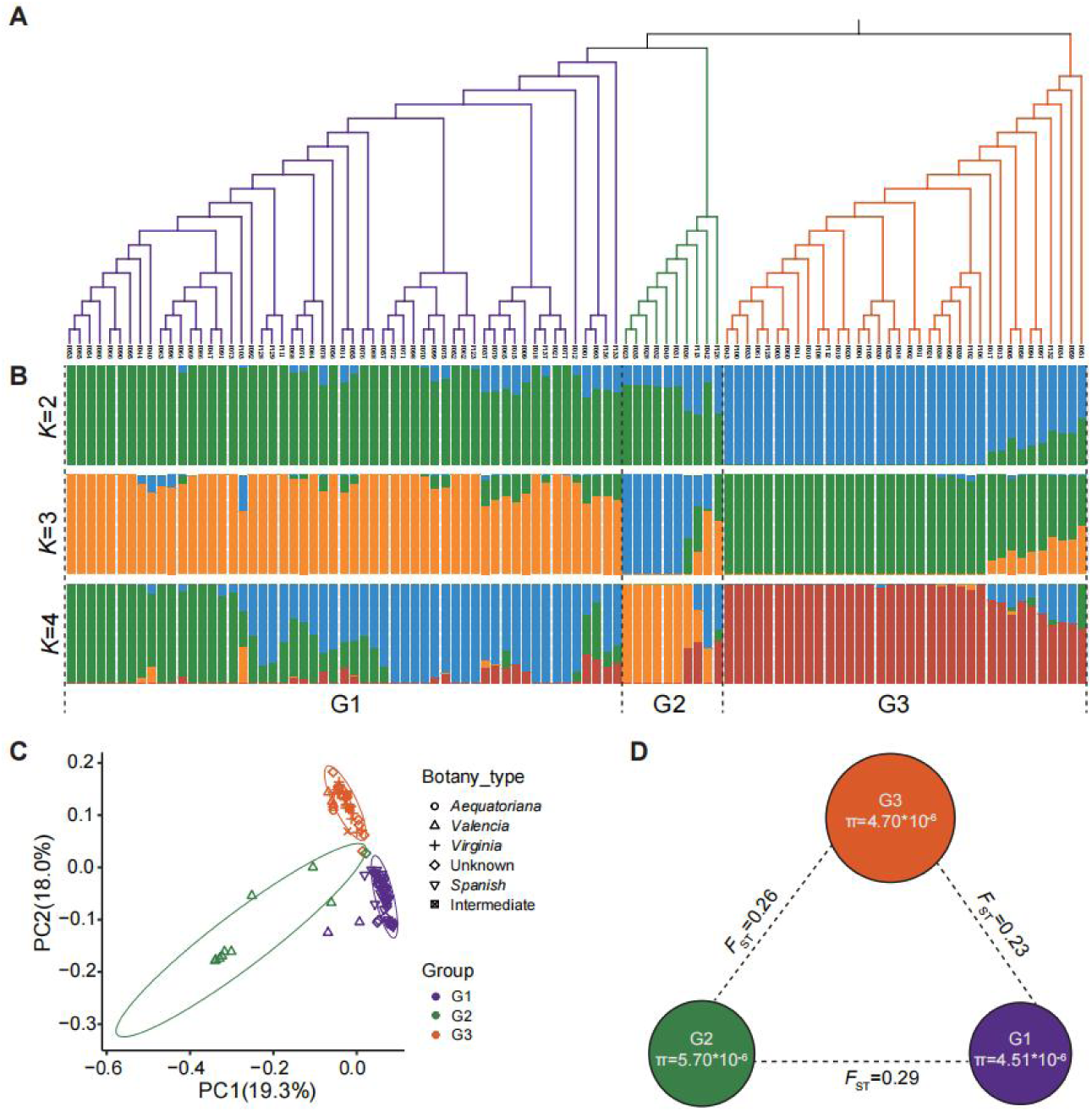
Population structure of 101 peanut accessions. **A** and **B**, phylogenetic tree and population structure analyses of all accessions. The structure analysis (K = 2 to 4) matches the phylogenetic tree. The color in the structure plot indicates the component state in each sample. **C**, principal component analysis (PCA) showing the first two principal components; dot colors correspond to the phylogenetic tree grouping. **D**, the diversity (π) and genetic distance (*F*st) across the groups, where color: phylogenetic group; radius of pie: genetic diversity; and dashed line length: *Fst* value between two subpopulations.

Based on the abovementioned phylogenetic and population structure analyses, we proceeded to calculate the fixation index (*F*_ST_) and nucleotide diversity (π) to assess the genetic divergence and diversity among the three subpopulations (Fig. 5 D). The *F*_ST_ values revealed that the greatest genetic distance was between G1 and G2 (0.29), followed by G2 and G3 (0.26), with the lowest *F*_ST_ values observed between G1 and G3 (0.23). This suggested that G1 and G2 have undergone more significant divergence compared to the other subpopulations, while accessions in G1 and G3 exhibit a relatively closer genetic affinity. Notably, the π values were highest in the G2 subpopulation (5.70×10^−6^), followed by G3 (4.70×10^−6^) and G1 (4.51×10^−6^), indicating a potentially higher gene flow between the G1 and G3 double-seed botanical types (Spanish and Virginia). Consequently, in future breeding initiatives, the genetic diversity of peanut cultivars could be broadened effectively by prioritizing genotypes within the G2 subpopulation.

## Discussion

The rapid advancements in high-throughput sequencing technology have led to a reduction in DNA sequencing costs by over 100,000 times [35], since the introduction of the first commercially available NGS instrument in 2008 [36]. Current cost estimates for widely used platforms such as the BGI DNBSEQ-T7 (~6.0 Tb data yield per day) and Illumina NovaSeq X series systems (~16.0 Tb data yield within 1–2 days) in China stand at approximately $1.43/Gb and $2.57/Gb, respectively. Despite the significant increase in sequencing throughput and cost efficiency, the expenses associated with library preparation procedures remain high, typically ranging from $14.28 to $28.57 per sample. This cost barrier particularly hampers breeders from utilizing NGS technologies for crops with gigabase-scale genomes, such as peanuts.

In this study, we presented a novel genotyping method dRAPD-seq, which involves only two easy-to-conduct steps to get a final DNA library: semi-specific PCR amplification and sequencing adapter ligation (Fig. 1). In contrast to conventional PCR which focuses on specific amplification using two forward and reverse primers [37], our approach employed a single AD primer (seen as a group of SP primers) to initiate semi-specific amplification under optimized cycling conditions. When two SP primers could bind randomly at an appropriate distance along the gDNA template during semi-specific PCR, the resulting amplicons could be detected by NGS technology. Therefore, dRAPD-seq utilized only one random AD primer sequence, although it is crucial to avoid AD primers consisting solely of NNNNNN bases based on our experience. When multiple samples are distinguished by unique barcodes tagged at a same AD primer and then PCR products from these samples could be pooled into one tube by AIO-seq method [32], the total library preparation cost per sample is nearly equivalent to that of a single PCR (~$1.75). Consequently, the dRAPD-seq genotyping method is cost-effective and easy-to-conduct for crop breeders and small-scale laboratories with limited budgets and equipment.

Compared to traditional low-throughput and gel-based molecular markers (for example, RAPD, RFLP and SSR), high-throughput genotyping technologies, especially WGS and GBS, have been extensively utilized in various aspects of plant population research, including germplasm evaluation [38,39], genetic mapping [40,41], biological evolution [5,9], etc. Despite the availability of many new NGS-enabled genotyping methods, the dRAPD-seq developed in this study presents several distinct advantages (Appendix K). First, unlike WGS and GBS, which require fragmentation of complete gDNA prior to library preparation [42,43], dRAPD-seq directly utilizes semi-specific PCR amplification of gDNA, achieving an equivalent fragmentation effect comparable to MRASeq [25], Hyper-seq [26], and target SNP-seq [24]. Second, different with GBTS [44], MRASeq and target SNP-seq, which require specific probes, programs or multiplex primers, dRAPD-seq only needs a single random AD primer to amplify partial genome sequences, thereby having the simplest primer design complexity among these methods. Third, by utilizing flexible barcodes attached to the AD primer and the AIO-seq strategy [32], dRAPD-seq incurs the lowest library preparation cost. Moreover, we have set up a fully automated workflow of dRAPD-seq experiment in our laboratory, which starts from gDNA of 96 samples (a batch), and ends with a final DNA library ready for NGS sequencing. While the genome coverage (~1–10%) and SNP count (~10^3^–10^4^) identified through dRAPD-seq are relatively lower (Supplementary Data Table S7), this method is better suited for applications in germplasm resource assessment, kinship inference, and gene mapping. Unlike genome-wide association studies (GWAS) that typically require hundreds of thousands (or even more) of SNP markers for higher resolution and accuracy [5,45], dRAPD-seq is more appropriate for these analyses, as hundreds to thousands of well-distributed markers are usually adequate [7,46,47].

After assessments of the dRAPD-seq genotyping method, we utilized it to explore the genetic relationships and population structure of a panel of peanut accessions, and observed a relatively low genetic diversity within this population. To our knowledge, cultivated peanut has undergone significant genetic bottlenecks during their evolutionary and domestication processes [48]. The general consensus suggests that cultivated peanuts likely originated from a singular hybridization event between two diploid progenitors, estimated to have occurred approximately 10,000 to 470,000 years ago [18,49]. The inherent self-pollination in cultivated peanuts would have led to reproductive isolation from their progenitor species [50], combined with limited seed dispersal capacity characteristic of peanuts, resulting in an initial narrow genetic foundation and restricted potential for genetic variation [48]. Previous studies on genetic diversity in another significant self-pollinating leguminous crop, soybean, have also shown low levels of genetic diversity. For instance, the average PIC and MAF of 277 Chinese and 300 American soybean accessions were reported as 0.26, 0.25, 0.24, and 0.22, respectively [51], comparable to our findings. In contrast, research on maize, a key cross-pollinating crop, revealed higher levels of genetic diversity, with the average PIC and genetic diversity of 344 Chinese maize breeding lines reported as 0.344 and 0.442, respectively [52]. These findings highlight that self-pollinating crops tend to exhibit lower genetic diversity compared to cross-pollinating crops. Furthermore, the selective pressures exerted through breeding practices may have further constrained the genetic diversity of cultivated peanuts. For example, an analysis of the genetic diversity within 32 peanut cultivars developed under the University of Georgia breeding program since its establishment in 1931 indicated a high degree of genetic similarity ranging from 94.6% to 99.9% [48]. Similarly, an examination of the genetic distance among 320 global peanut collections ranged from 0.043 to 0.127, with an average of 0.080 [10]. These outcomes are in line with our observations and collectively underscore the imperative of broadening the genetic base of cultivated peanuts in future breeding endeavors.

Population structure analysis is crucial for understanding the genetic background and relationships among diverse germplasms, providing valuable insights for preliminary genetic resource assessment. In our investigation, phylogenetic, population structure, and principal component analyses categorized the 101 peanut accessions into three main subpopulations representing Spanish, Valencia, and Virginia types, respectively. Correspondingly, Zheng et al. [10], Brown et al. [48], Hsu et al. [2] and Liu et al. [9] employed similar classification methods for 320 global peanut collections, 31 Taiwan province peanut germplasms, 32 peanut cultivars from the University of Georgia, and 203 global peanut cultivars, respectively, based on their botanical types. We also found that certain genome-wide association studies (GWAS) have delineated peanut collections into subsp. *fastigiata* and subsp. *hypogaea*, encompassing distinct botanical types [6,53,54]. In our study, we observed that Spanish and Valencia types clustered into separate subpopulations due to the predominance of Chinese landraces in the Valencia type and modern cultivars in the Spanish type, with limited genetic exchange occurring between the two types in China. This phenomenon can also be observed in African core groundnut collection [55]. Moreover, *F*_ST_ and π values indicated that the Spanish and Virginia subpopulations exhibited closer genetic proximity and lower nucleotide diversity compared to the Valencia subpopulation. This finding aligns with a prior analysis of 203 global peanut cultivars, demonstrating that Valencia types displayed greater genetic distance and higher nucleotide diversity than Spanish and Virginia types, with Virginia types showing the lowest nucleotide diversity [9]. In China, the double-seed botanical varieties (e.g. Spanish and Virginia) are considered to exhibit higher productivity compared to multiple-seed botanical types (e.g. Valencia). Spanish and Virginia varieties prevail as the primary peanut cultivars in the southern and northern regions, respectively, with frequent genetic exchanges observed between these varieties as well as between the northern and southern regions. For example, certain Virginia type accessions (P002 (‘Huayu 910’), P013 (‘Huayu51’), and P030 (‘Jihua22’); Table S1) exhibit Spanish type ancestry, tracing back to the renowned Chinese landrace ‘Fuhuasheng’ (http://peanut.cropdb.cn/variety/). These inter-type genetic exchanges manifest in closer genetic distances and lower nucleotide diversity. Therefore, diversifying the use of other peanut types may be a key strategy to broaden the genetic foundation of cultivated peanuts.

In conclusion, we have presented dRAPD-seq, an innovative whole-genome genotyping method that utilizes semi-specific PCR amplification by AD primer and sequencing, which is well-suited for diverse applications in plant genetics and molecular breeding efforts. Furthermore, the genetic relationships and population structure revealed within the peanut germplasm population could provide valuable insights for the strategic utilization of these accessions in peanut breeding programs.

## Materials and Methods

### Plant Materials and DNA Isolation

A total of 101 peanut accessions were included in this study, comprising 90 domestic and 11 foreign accessions (Supplementary Data Table S3). Prior to DNA extraction, all peanut accessions and a standard rice cultivar Nipponbare (*Oryza sativa* L.) were cultivated under greenhouse conditions at the Zhanjiang Experiment Station, Chinese Academy of Tropical Agricultural Sciences. For each sample, young leaves were snap-frozen in liquid nitrogen and stored at −80◦C. Total gDNA was extracted using a CTAB-based method [56]. The quality of the gDNA was first assessed by 0.8% agarose gel electrophoresis and then the quantity was measured using NanoDrop 2000 (Thermo Fisher Scientific) and Qubit™ 4.0 fluorometer (Invitrogen). The gDNA with no visible degradation was used for subsequent library preparation.

### Library Preparation and Sequencing

Semi-specific PCR amplification was conducted as follows. The 50-μL PCR mixtures comprised 30 ng of gDNA, 10 μL 5× TAB buffer, 20 μL of AD primer (10 μM), and 1 μL of TAE polymerase (TruePrep® DNA Library Prep Kit V2 for Illumina®, Cat. no. TD501-02, Vazyme Biotech). PCR thermocycling was carried out in Biometra Tone 96G thermocyclers (Analytik Jena), with the following conditions: Choose the preheat lid option and set to 105°C; pre-denaturation at 98°C for 2 min; followed by five cycles of 98°C for 30 s, 30°C for 60 s, ramping to 72°C (0.3°C/s) and 72°C for 3 min; then 20 cycles of 98°C for 30 s, 40°C for 30 s, 72°C for 25 s; and a final extension at 72°C for 10 min. For 101 peanut accessions, 24 AD primers (each ligated with a unique barcode at 5’-end, Supplementary Data Table S1) were initially designed for the amplification of gDNA from each of 24 peanut accessions, respectively. The fragment size profile of each sample’s PCR product was analyzed by Qsep 100 bioanalyzer (BiOptic). Subsequently, the amplified products of 24 samples were pooled into one tube (a group) by AIO-seq pooling strategy [32]. After that, TruSeq libraries was prepared for each group according to the manufacturer’s instructions for Hieff NGS® 384 CDI Primer for Illumina® (Cat. 12412ES02, Yeasen Biotech) and Hieff NGS® Ultima Pro DNA Library Prep Kit for Illumina® (Cat. 12201ES96, Yeasen Biotech). Within each resulting TruSeq library, fragments within the peak size range of approximately 400–650 bp were isolated using the cassette-based Pippin HT system (Sage Science). Finally, these selected fragments were sent to Berry Genomics Company (Beijing, China) and sequenced on an Illumina NovaSeq 6000 system utilizing a paired-end 150 bp (PE150) sequencing strategy.

To evaluate the accuracy of SNP genotype amplified using the AD primer, gDNA from the identical rice Nipponbare line was initially fragmented utilizing the Covaris S220 ultrasonic DNA breaker (Covaris). Subsequently, the fragmented DNA were subjected to whole-genome sequencing (WGS) library preparation employing the VAHTS Universal DNA Library Prep Kit for Illumina V3 (Cat. ND607-01, Vazyme Biotech) following the recommended protocols. Lastly, the prepared library was sent to Berry Genomics Company (Beijing, China) for PE150 sequencing.

### Read Mapping, tag identification and SNP calling

For raw WGS reads of rice Nipponbare, low-quality reads were filtered out using fastp (version 0.23.0) [57], and then aligned to the R498 reference genomes [34] using BWA-MEM (version 0.7.17) [58]. The VCF file was produced by HaplotypeCaller module (parameters: --min-base-quality-score 20 --minimum-mapping-quality 30) implemented in Genome Analysis Toolkit (GATK; version 4.2.5.0) [59], and then SNP variations were selected by SelectVariants module. After that, origin SNPs were filtered by VariantFiltration module with the following parameters: QD < 2.0 || MQ < 40.0 || FS > 60.0 || SOR > 5.0 || MQRankSum < −12.5 || ReadPosRankSum < −8.0. Finally, only SNPs with the sequencing depth of at least 5× were retained and designated as the gold standard.

For raw dRAPD-seq reads of rice Nipponbare, the barcode sequences were trimmed off from Read1 and Read2 using fastp (version 0.23.0) [57], and then aligned to the Nipponbare_V7.0 reference genome [33] using BWA-MEM (version 0.7.17) [58]. SAMtools software (version 1.9) [60] was used to calculate the depth of each base by utilizing properly aligned read pairs and alignments with high mapping quality (MQ ≥ 30). Similar to a previous study [27], genomic loci meeting the following criteria were designated as a sequence tag: (i) The number of mapped Read1 or Read2 starting at a specific genomic coordinate should be at least three, and (ii) the depth of a tag position should be at least 3× higher than that of the previous base (where the coordinate of the previous base corresponds to the tag position −1 when Read1/Read2 is aligned to the positive strand of the reference genome or tag position +1 when aligned to the negative strand). The visualization of genome-wide tag distribution was achieved by an R package CMplot (version 4.2.0) [61].

For peanut accessions, the raw dRAPD-seq sequencing reads of a group (24 accessions) were first de-multiplexed into corresponding sample by fastq-multx tool (version 1.4.3) [62] according to the unique barcode sequences present at the start of Read1 and Read2. After splitting, raw reads were trimmed by fastp (version 0.23.0) [57], and then aligned to the peanut Tifrunner.gnm2.J5K5 reference genome [1] using BWA-MEM (version 0.7.17) [58]. As mentioned above, only high-quality alignments were retained and used to calculate the genome coverage by Qualimap2 (version 2.2.1) [63]. Bam files were sorted by SAMtools (version 1.9) [60] and Read group information was added to each sample’s alignment files by Picard AddOrReplaceReadGroups module (version 2.26.7; http://broadinstitute.github.io/picard/). Individual GVCF file of each sample was generated by HaplotypeCaller module (parameters: --min-base-quality-score 20 --emit-ref-confidence GVCF) and then all GVCF files were merged into an integrated file by CombineGVCFs module [59]. Further, GenotypeGVCFs and SelectVariants modules were applied to detect variations and get a raw VCF set comprising only SNPs. VariantFiltration module was used to remove the low-quality SNPs with the following parameters: QD < 2.0 || MQ < 40.0 || MQRankSum < −12.5 || ReadPosRankSum < −8.0 || FS > 60.0. Finally, VCFtools (version 0.1.12b; parameters: --max-alleles 2 --min-alleles 2 --minQ 30 --max-missing 0.8 --maf 0.05) [64] was utilized to select a subset of qualified SNPs for downstream analyses.

### Genetic diversity and population structure analysis

Various genetic diversity metrics were calculated based on the qualified SNP data. The polymorphic information content (PIC) was determined by PIC_CALC software (version 0.6; https://github.com/luansheng/PIC_CALC). Minor allele frequency (MAF), observed heterozygosity (*Ho*) and genetic distance were obtained by Plink software (version 1.9) [65]. Simpson’s index of diversity was assessed using a custom Perl script according to the method outlined in a previous study [66].

To perform phylogenetic analysis, the filtered VCF file was converted into a PHYLIP file using vcf2phylip Python program [67]. Subsequently, the PHYLIP file was imported into MEGA11 (version 11.0.13) [68] to generate a neighbor-joining (NJ) tree with the bootstrap test of 1,000 replicates. The final tree topology was visualized using the online tool iTOL (version 6.6; https://itol.embl.de/). The genetic structure and estimated admixture proportions were investigated using the ADMIXTURE software (version 1.3.0) [69]. The number of genetic clades was predefined from K = 1 to 10. The ancestry distributions of accessions were visualized using R script. Principal component analysis (PCA) was performed using PLINK software (version 1.9) [65], and the first two eigenvectors obtained from PCA were plotted in two dimensions. Similar to previous studies [5,9], pairwise fixation index (*F*_ST_) and nucleotide diversity (π) were computed in the VCFtools program (version 0.1.12b) [64] with a 50-kb non-overlapping sliding window along each chromosome. The π and *F*_ST_ between different subpopulations were calculated based on the average value of all 50-kb sliding windows.

## Acknowledgements

This work was supported by the National Natural Science Foundation of China (32300340), the R&D program of Shenzhen (KCXFZ20211020164207012), Hainan Provincial Natural Science Foundation of China (324QN302, 321QN348), and Central public-interest Scientific Institution Basal Research Fund (1630102024002).

## Author contributions

Z.X., Y.C., and S.Z. discussed and designed the experiments; S.Z., Z.X., and Y.C. wrote the manuscript with contributions from all authors; S.Z., Y.W., X.Z., and S.X. performed the experiments; S.Z., Y.W., H.C., Y.Y., J.G., and P.C. participated in the data analysis; S.Z., Z.X., and Y.W. prepared the figures; Z.X. and Y.C. supervised the project. All authors have read and approved the manuscript.

## Data Availability

The raw sequence data reported in this paper have been deposited in the Genome Sequence Archive in the National Genomics Data Center under accession number CRA015526 and are publicly accessible at https://ngdc.cncb.ac.cn/gsa.

## Conflict of interests

The authors declare that they have no conflict of interest. S.Z. and Y.C. have filed a Chinese patent application CN202211418813.3 based on this work.

## Abbreviations

AD: arbitrary degenerate primer
AFLP: amplified fragment length polymorphism
AIO-seq: all-in-one sequencing
BWA: Burrows-Wheeler Aligner
Chr: chromosome
dRAPD-seq: degenerate random amplification polymorphic DNA and sequencing
FBI-seq: foreground and background integrated genotyping by sequencing
*F*_ST_: fixation statistics
GATK: Genome Analysis Toolkit
Gb: Gigabases
GBS: Genotyping-by-Sequencing
GBTS: genotyping by target sequencing
gDNA: genomic DNA
GWAS: genome-wide association studies
*Ho*: observed heterozygosity
IGV: Integrative Genomics Viewer
MAF: minor allele frequency
Mb: Megabases
MQ: mapping quality
MRASeq: multiplex restriction amplicon sequencing
NGS: next-generation sequencing
NJ: neighbor-joining
π: nucleotide diversity
PCA: principal component analysis
PE150: paired-end 150 bp
PIC: polymorphic information content
PTMA: primer-template mismatched annealing
PTPA: primer-template perfect annealing
QTL: quantitative trait locus
QUAL: Phred-scaled quality score
RAPD: random amplification polymorphic DNA
RFLP: restriction fragment length polymorphism
SNP: single nucleotide polymorphism
SSR: simple sequence repeat
SP: specific primer
WGG: whole-genome genotyping
WGS: whole-genome sequencing.

## Supplemental information

**Table S1** Arbitrary degenerate primer sequences.

**Table S2** Basic statistics of sequencing reads and tags amplified by AD primer.

**Table S3** Sequencing data information and statistics of 101 peanut accessions.

**Table S4** SNP number distributed on peanut genome.

**Table S5** Summary statistics of 1203 high-quality SNP markers.

**Table S6** Genetic distance matrices among 101 peanut germplasms.

**Table S7** Comparison between dRAPD-seq and other NGS-based genotyping methods.

## Supplemental Figures

**Figure S1.**
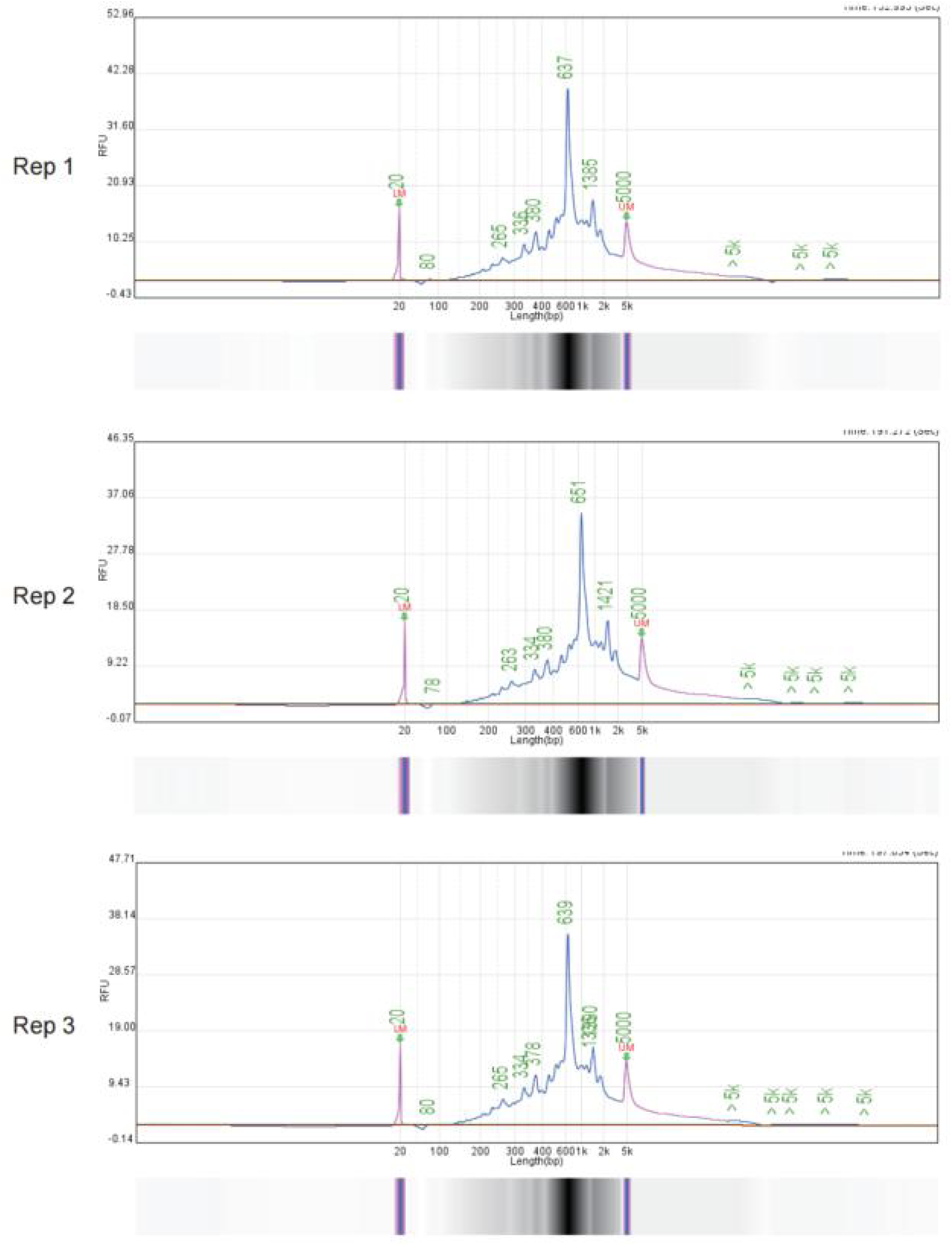
Size profile of PCR products for three technical replicates. Fragment size distribution was measured by Qsep 100 bioanalyzer.

**Figure S2.**
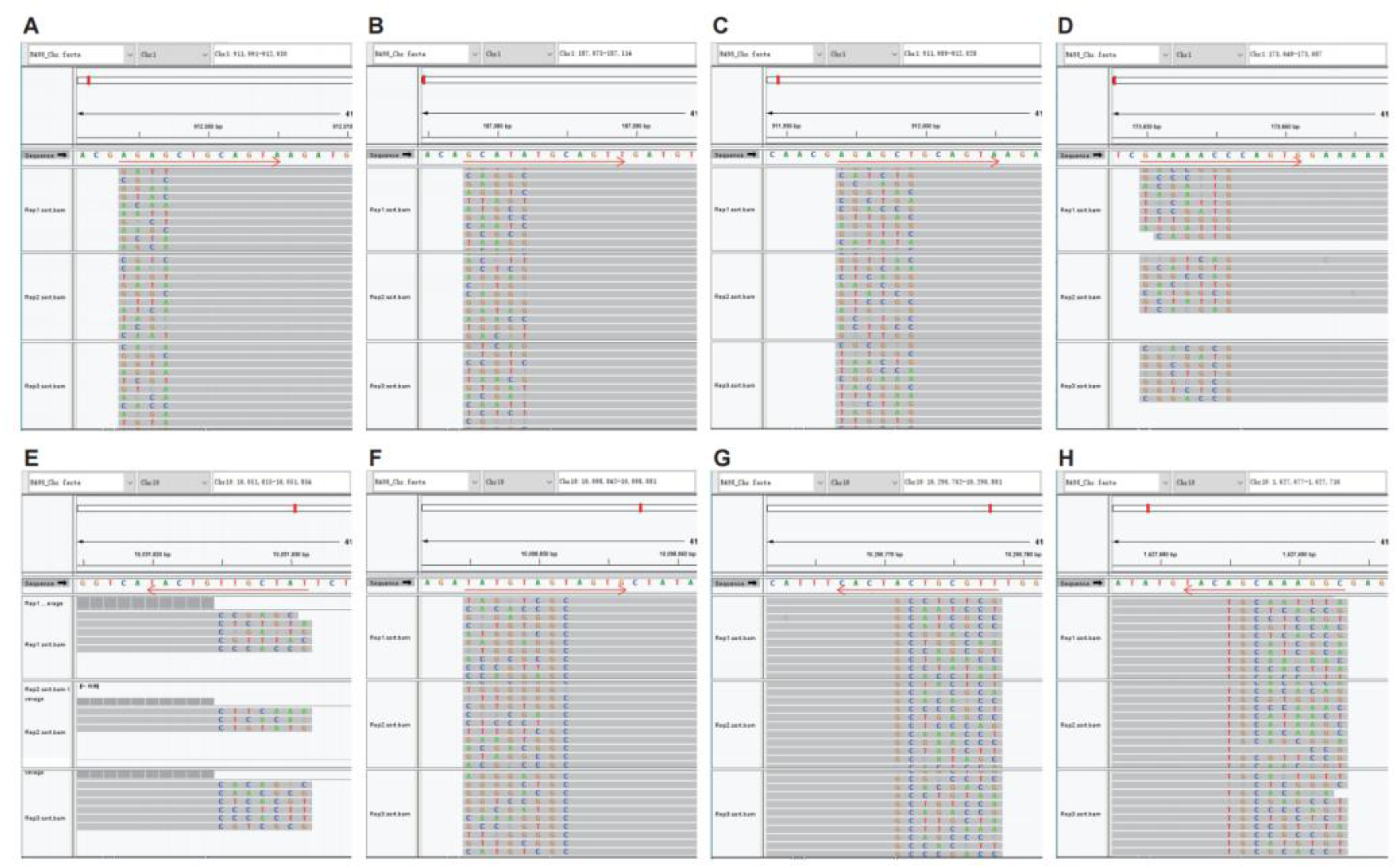
Snapshots of the Integrative Genomics Viewer (IGV) browser displaying that AD primer was bound with DNA template by primer-template mismatched annealing (PTMA) and dRAPD-seq reads were accumulated to form a tag. Red left/right arrow represented the binding site of AD primer. A-D, tags formed by 4-nt, 5-nt, 6-nt and 7-nt soft-clipped reads, respectively. E-H, tags formed by 7-nt, 8-nt, 8-nt and 9-nt soft-clipped reads, respectively.

**Figure S3.**
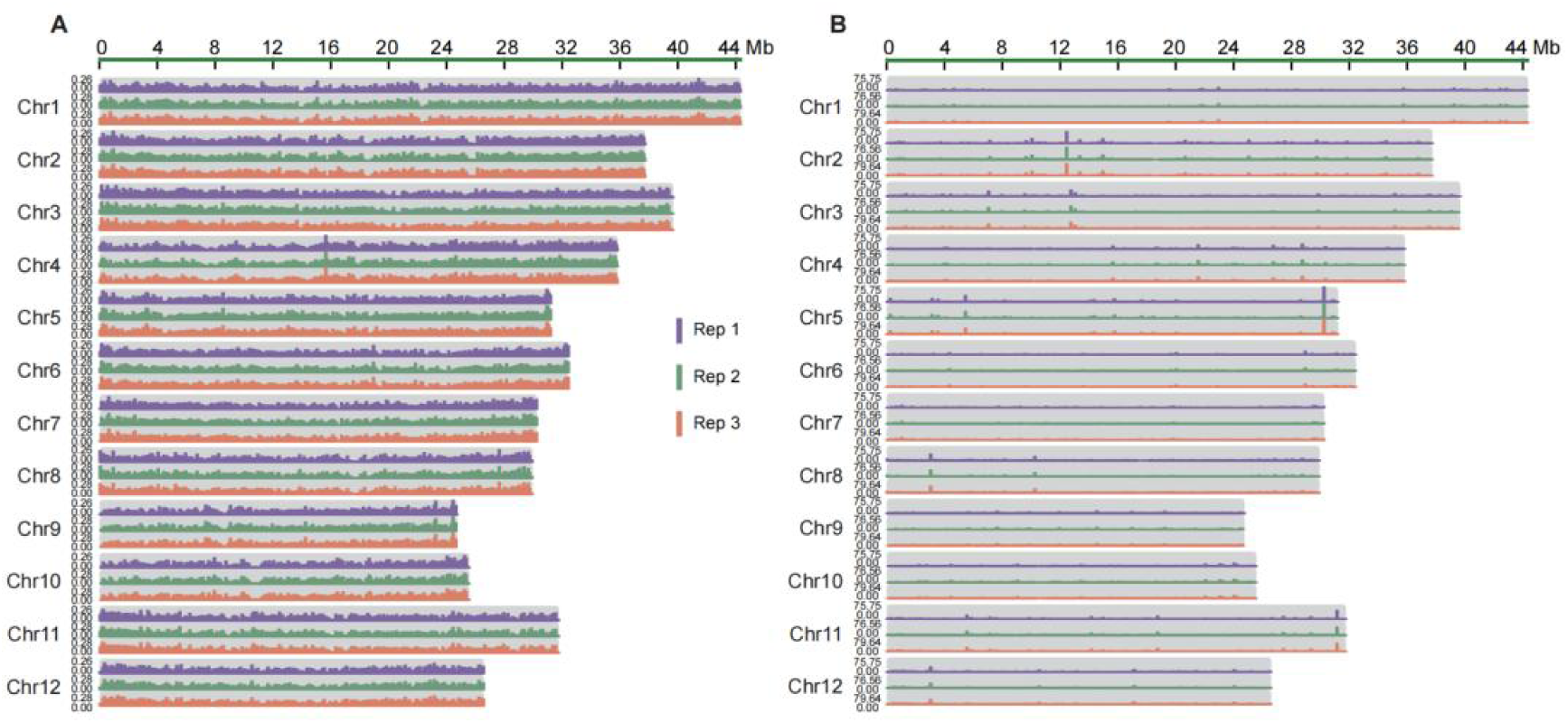
Genome coverage (A) and mean depth (B) of dRAPD-seq reads across the rice genome calculated within a 0.1-Mb bin size. Totally 1.0 Gb of sequencing data were randomly subsampled from each of three replicates for analysis.

**Figure S4.**
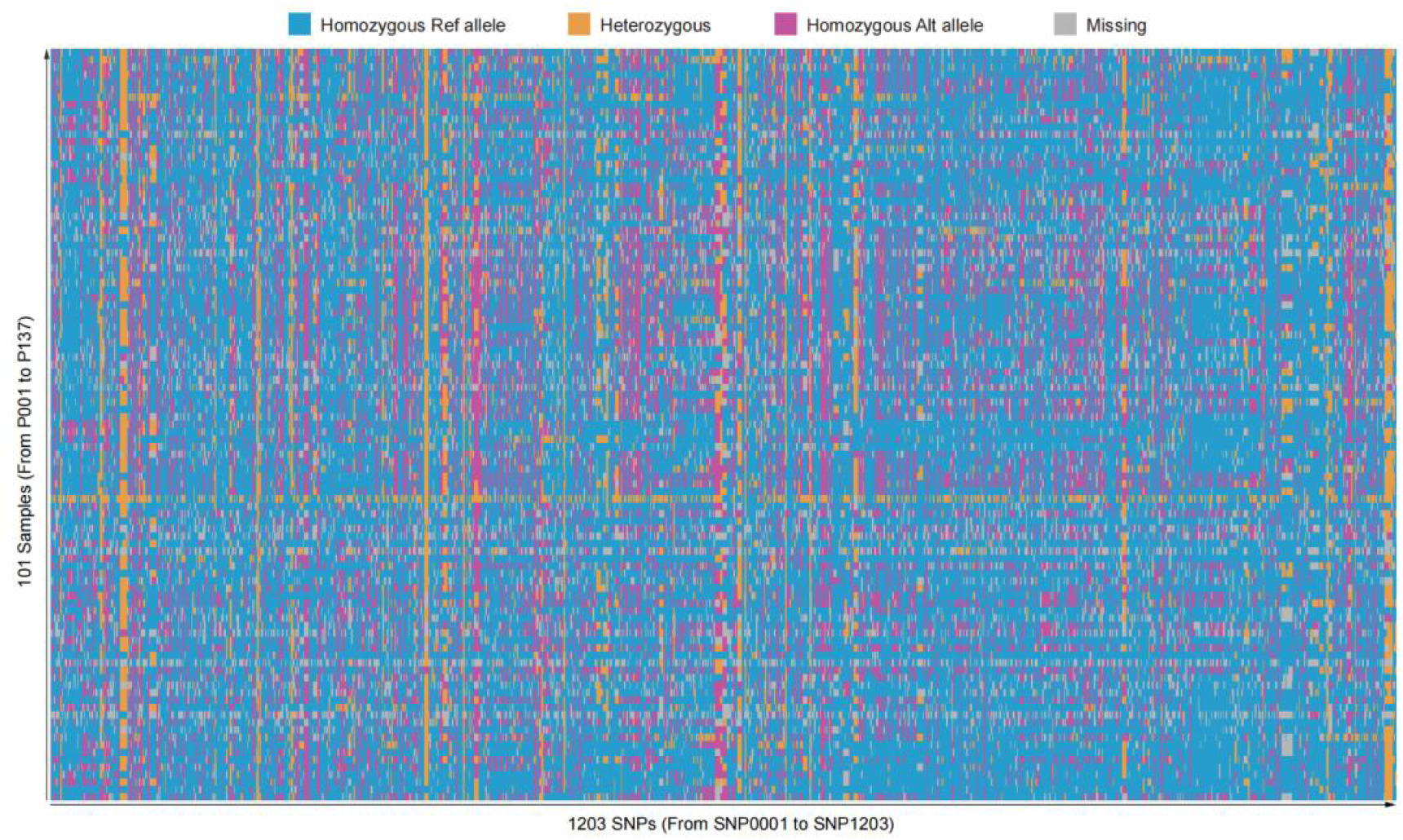
Graphical representation of DNA fingerprint for 101 peanut accessions using 1203 SNP markers.

## Notes

### Competing Interest Statement

The authors have declared no competing interest.

